# The beetle amnion and serosa functionally interact as apposed epithelia

**DOI:** 10.1101/025247

**Authors:** Stefan Koelzer, Maarten Hilbrant, Thorsten Horn, Kristen A. Panfilio

## Abstract

Unlike passive rupture of the human chorioamnion at birth, the insect extraembryonic (EE) tissues – the amnion and serosa – actively rupture and withdraw in late embryogenesis. Despite its importance for successful development, EE morphogenesis remains poorly understood. Contradicting the hypothesis of a single, fused EE membrane, we show that both tissues persist as discrete epithelia within a bilayer, using new tissue-specific EGFP transgenic lines in the beetle *Tribolium castaneum*. Quantitative live imaging analyses show that the amnion initiates EE rupture in a specialized anterior-ventral cap, while RNAi manipulation of EE tissue complement and function reveals that the serosa is autonomously contractile. Thus the bilayer efficiently coordinates the amnion as initiator and serosa as driver to achieve withdrawal. The novel bilayer architecture may reflect evolutionary changes in the EE tissues specific to holometabolous insects. More generally, tissue apposition in a bilayer exemplifies a high degree of functional interaction between developing epithelia.

## INTRODUCTION

Embryogenesis requires dynamic interaction between tissues to create changing three-dimensional configurations, culminating in the completion of the body. In parallel to the amniote vertebrates (Calvin and Oyen, 2007), the insects have evolved extraembryonic (EE) tissues that arise in early embryogenesis to envelop the embryo (Panfilio, 2008). Like its vertebrate namesake, the amnion encloses a fluid-filled cavity around the embryo. As the outermost cellular layer, the serosa provides mechanical and physiological protection (Rezende et al., 2008; Jacobs et al., 2013; Jacobs et al., 2014). This protective configuration is not permanent, though, and a major reorganization of the EE tissues, including final apoptosis, is essential for embryos to correctly close their backs in late development. For these events, perhaps the nearest morphogenetic equivalent in the model system *Drosophila* is the retraction of the wing imaginal disc during metamorphosis, where the squamous peripodial epithelium also everts, contracts, and undergoes apoptosis (Aldaz et al., 2010). However, research on *Drosophila* cannot address the morphogenesis of the two EE epithelia directly, due to the secondarily derived nature of the single EE tissue, the amnioserosa, which does not surround the embryo (Schmidt-Ott, 2000; Rafiqi et al., 2012).

The overall process of insect EE withdrawal – the active process whereby the EE tissues withdraw from the embryo and leave it uncovered – has been addressed in classical descriptions for many species (reviewed in Panfilio, 2008). However, a major open question has been the organization and role of the amnion. This is primarily because it is difficult to visualize in its native topography with respect to other tissues, due to a lack of amnion-specific molecular markers (discussed in Koelzer et al., 2014) and to the histological similarity of the two mature EE tissues as simple, squamous epithelia (Panfilio and Roth, 2010).

Here, we present the first clear determination of the relative topography and role of the amnion in late development in a holometabolous insect, the red flour beetle, *Tribolium castaneum*. We characterize an enhancer trap line that labels the amnion and use this in conjunction with a recently characterized serosal line (Koelzer et al., 2014) to morphogenetically dissect which tissue is responsible for which aspects of EE tissue withdrawal. The topographical arrangement of the tissues differs strikingly from what was previously known in hemimetabolous insects and what had previously been hypothesized for *Tribolium*, which has consequences for understanding the entire withdrawal process. Furthermore, we provide evidence that while the serosa strongly drives the contraction and folding of the tissues, the amnion initiates EE rupture.

## RESULTS AND DISCUSSION

### The amnion and serosa form a persistent bilayer during late development

To augment the toolkit for tissue-specific visualization in *Tribolium*, we identified and characterized an enhancer trap line with amniotic EGFP expression (Fig. 1, see also Figure 1-figure supplement 1, Movie S1). Prior to withdrawal morphogenesis, the EGFP-labeled tissue fully envelops the embryo but does not cover the yolk (Fig. 1A-B). To confirm that this tissue is indeed the amnion, and not a specialized region of the serosa, we examined EGFP expression after RNAi for *Tc-zen1*, thereby eliminating serosal tissue identity (van der Zee et al., 2005). In the absence of the serosa, the amnion occupies a dorsal position over the yolk, and indeed this tissue expresses EGFP (Fig. 1C-D).

**Figure 1.**
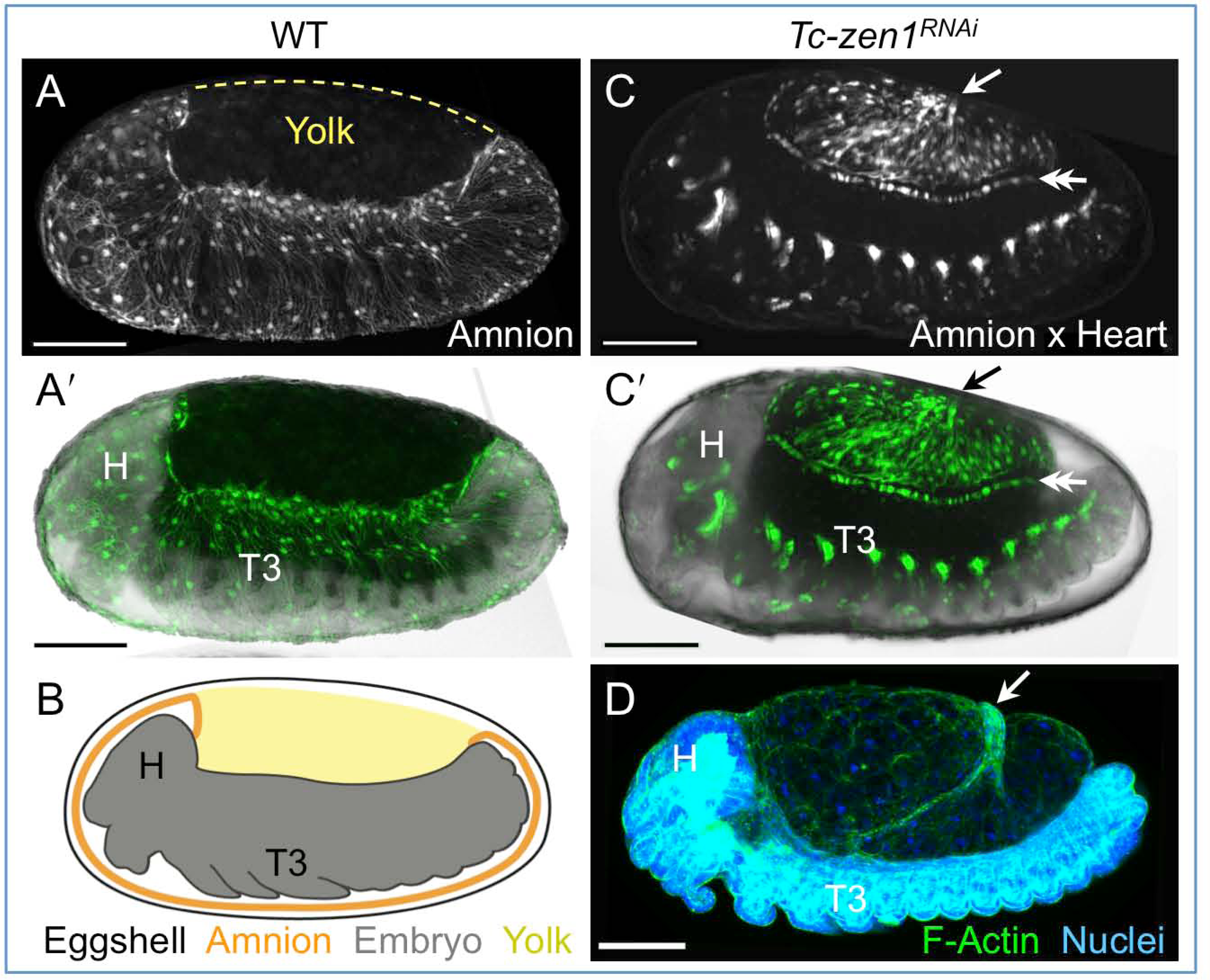
The enhancer trap line HC079 is an autonomous amniotic tissue marker. Images are lateral, with anterior left and dorsal up, shown as z-stack maximum intensity projections or mid-sagittal schematic. Visualization reagents are indicated. **A-B.** In WT, EGFP expression is extraembryonic (EE) in a ventral domain that fully covers the embryo but not the yolk. See also Figure 1-figure supplement 1, Movie S1. **C-D.** Consistent with the WT EGFP domain being amniotic, the entire EE tissue expresses EGFP when serosal identity is eliminated after *Tc-zen1^RNAi^*. Here, the EE tissue does cover the yolk (dorsal to the cardioblast cell row: double-headed arrow), and acquires a characteristic ‘crease’ (arrow) during withdrawal morphogenesis (Panfilio et al., 2013). Scale bars are 100 *μ*m. Abbreviations: H, head; T3, third thoracic segment.

We then used the imaging lines to address the arrangement of the amnion and serosa during late development. Initially the two EE tissues are physically separate (Handel et al., 2005), but they progressively come together during early germband retraction until the only visibly distinct amniotic region is a rim of tissue at the embryo’s dorsal margin (Fig. 2A-A2). Previous histological studies in *Tribolium* and other holometabolous insects concluded that the region of overlap comprised a single EE cell layer, the EE tissues having intercalated or otherwise “fused” (*e.g*., Patten, 1884; Kobayashi and Ando, 1990; van der Zee et al., 2005). Alternatively, this structure was interpreted as only serosa (Panfilio et al., 2013), as the underlying amniotic region undergoes apoptosis in a hemimetabolous insect, the bug *Oncopeltus fasciatus* (Panfilio and Roth, 2010). In fact, we find that both tissues persist as apposed and very thin but distinct squamous epithelial layers (Fig 2B-D, 5C, Figure 2-figure supplement 1). This apposition is maintained throughout withdrawal morphogenesis (Figs. 2E-J, 5D). The new amnion-EGFP line thus sheds light on EE tissue structure during withdrawal: rather than a single, pseudostratified tissue, each EE tissue persists as a monolayer. Not only is this a novel EE tissue organization compared to what is known for other species, as we show below, this organization has several implications for how withdrawal morphogenesis proceeds.

**Figure 2.**
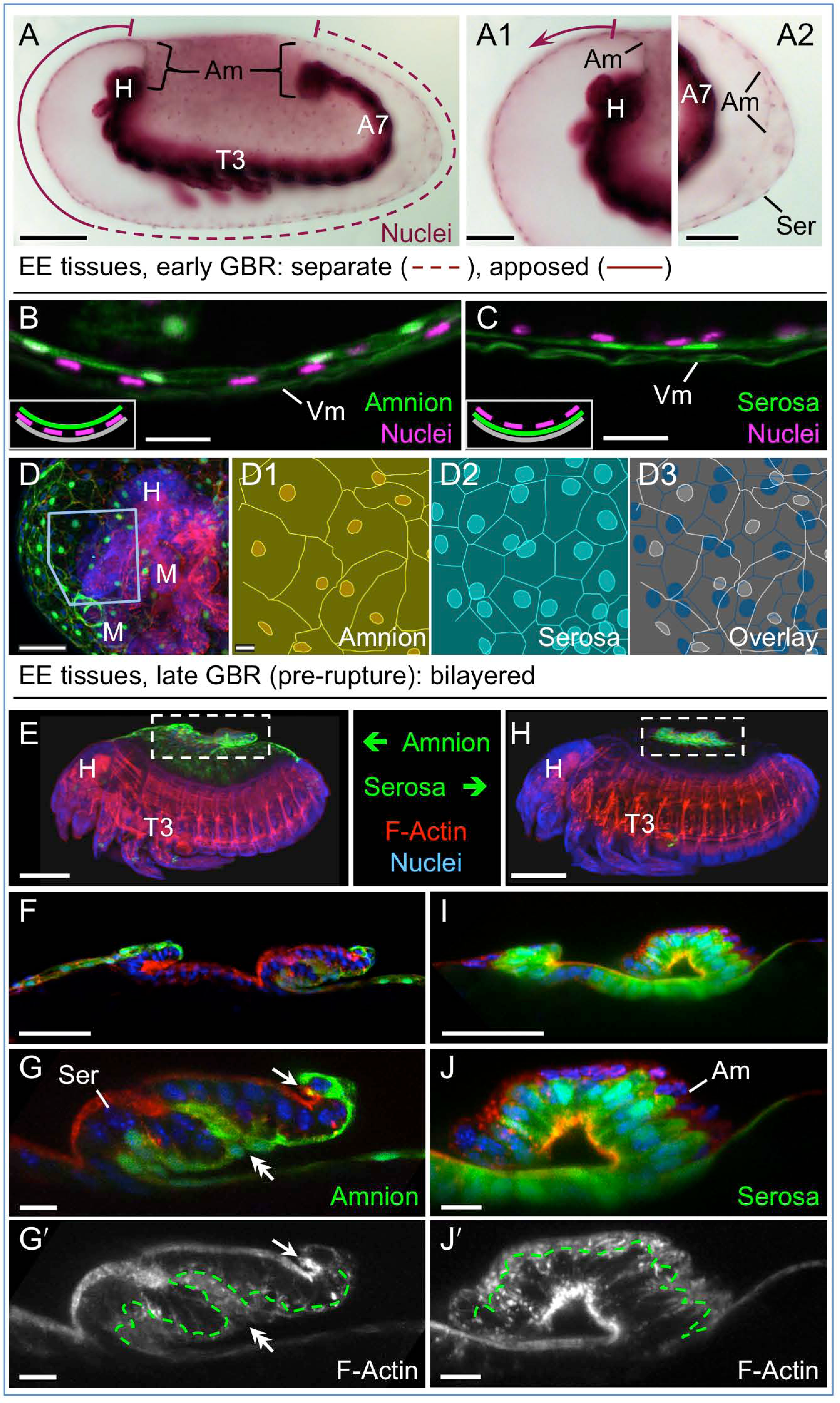
The amnion and serosa form a bilayer that moves as a single unit during late extraembryonic morphogenesis. Images are lateral, with anterior left and dorsal up, shown as sagittal optical sections (A-C,F,G,I,J) or maximum intensity projections (D,E,H). **A.** During germband retraction (GBR), the amnion (Am) progressively comes together with the serosa (Ser), shown at an intermediate stage where the tissues are apposed anteriorly (A1) but not posteriorly (A2). The preparation method results in embryo shrinkage (Wigand et al., 1998), amplifying apparent amniotic cavity volume, but without altering tissue topography, which is consistent across dozens of stage-matched specimens. **B-D.** Prior to rupture, the ventral EE tissue subjacent to the eggshell (autofluorescent vitelline membrane, Vm) is in fact comprised of distinct serosal (outer) and amniotic (inner) layers. Viewed in sagittal sections (B,C), tissue-specific EGFP labels continuous tissue sheets (partially masking the nuclear stain), while nuclei of the apposed EE tissue remain EGFP-negative and in a separate layer (see inset). Projections specifically in the anterior-ventral region also show two epithelial layers, which can be distinguished by tissue-specific cellular morphologies (see also Figure 2-figure supplement 1). **E-J.** Bilayered structure during early serosal compaction. Annotations: arrow, edge of both tissues; double-headed arrow, double bend in the tissues; dashed green lines delimit EGFP domains. Scale bars are 100 *μ*m (A,E,H), 50 *μ*m (A1,A2,D,F,I) and 10 *μ*m (B,C,D1-D3,G,G′,J,J′). Abbreviations: A7, seventh abdominal segment; M, mandible; and as defined above and in Fig. 1.

### The amnion initiates EE withdrawal

Given that the amnion persists as a discrete tissue throughout withdrawal morphogenesis, we then investigated its role in this process. Based on three lines of evidence, we conclude that the amnion initiates EE rupture at the onset of withdrawal.

Firstly, close inspection of the amnion EGFP signal shows that an anterior-ventral cap of cells are morphologically distinct: while most amniotic cells are striated and elongated along the dorsal-ventral axis, cells within this region are rounder and have more distinct cell outlines and brighter EGFP signal (Fig. 3A, see also Figure 1-figure supplement 1, Figure 2-figure supplement 1), and rupture begins within this territory (Fig. 3B, see also Movie S2).

**Figure. 3.**
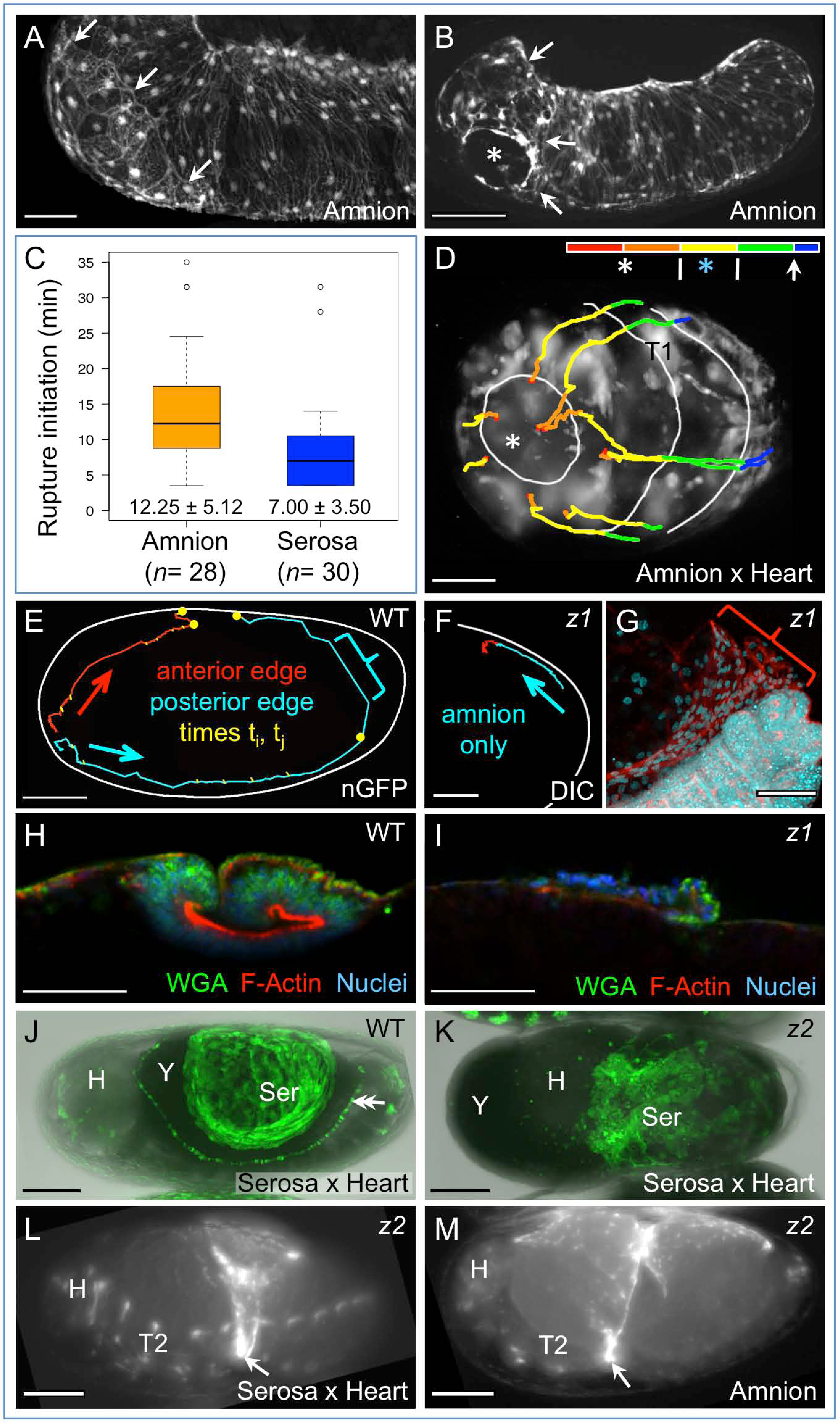
Rupture dynamics: amniotic initiation and serosa-driven progression. Images are lateral with anterior left and dorsal up, unless ventral (D,K) or dorsal-lateral (J), shown as maximum intensity projections (A,B,D,G,J,K) or sagittal optical sections (H,I). **A-B.** Amnion-EGFP signal before (A) and during (B) rupture: arrows delimit the anterior-ventral cap of morphologically distinct cells; the asterisk marks the tissue opening (see also Movie 2). **C.** Box plot showing that rupture initiation, as defined in the Methods section, is longer in the amnion (values are median ± median absolute deviation). **D.** Tissue opening in a representative embryo filmed at high temporal resolution (see Methods). The colored time scale shows duration of track segments over 36.7 minutes at 19.5±1°C. Along the time scale and superimposed on the embryo are the site of rupture (white asterisk) and the withdrawing EE tissue edge (white lines, line with arrowhead marks time point shown). Tracks are shown for selected nuclei. For comparison, rupture is also indicated along the time scale for a morphologically stage matched embryo with serosal GFP (blue asterisk; see also Figure 3-figure supplement 1, Movie 3). **E-I.** Tracking and histological staining of withdrawing EE tissue edges. The WT EE tissue edge squeezes, then rapidly clears, the abdomen (E: blue bracket: 1-minute interval, 6.7-hour total track, 21°C). The distance of the EE track from the vitelline membrane (white line) reflects the degree of squeezing of the abdomen. After *Tc-zen1*^*RNAi*^ (*z1*), an amnion-only posterior edge contracts slowly over an uncompressed abdomen (F: 3.9-hour total track, 21°C), with ruffling of the tissue (F: red track segment; G: red bracket, same staining reagents as in H) and no F-actin enrichment in the folding tissue (I, compare with H and Fig. 2I-J). **J-K.** While WT serosal contraction leads to a dorsally condensed tissue (J), in strong *Tc-zen2*^*RNAi*^ (*z2*) phenotypes, the serosa condenses ventrally over the unopened amnion and the confined embryo (K). The double arrowhead labels the cardioblasts; “Y” marks the opaque yolk. **L-M.** In weaker *Tc-zen2*^*RNAi*^ (*z2*) phenotypes the EE tissues can rupture and tear ectopically, leaving a “belt” of EE tissue (arrows) that squeezes the embryo (dorsal-lateral views). Scale bars are 50 *μ*m (A,D,H,F-I) and 100 *μ*m (B,E,J-M). Panel A shows the same embryo as in Fig. 1A.

Secondly, rupture initiation is significantly longer in the amnion than in the serosa (Fig. 3C-D; *p*= 0.0005002, Mann-Whitney U test; see Methods), even when slight differences in developmental rate between the lines are considered (see also Figure 1-figure supplement 1). This is consistent with rupture being initiated by the amnion and a slight delay before the serosa also participates. Indeed, the difference in initiation duration corresponds morphologically to only a slight opening of the amnion, within the anterior-ventral cap region (82%, *n*= 28), before the serosa would also perceptibly rupture (see also Figure 3-figure supplement 1, Movie S3). We therefore hypothesize that the amniotic anterior-ventral cap cells represent a rupture competence zone and may differ from the rest of the tissue in the nature of its attachment to the serosa (Figure 2-figure supplement 1D-E), given that the edges of the two tissues then retain apposition throughout subsequent withdrawal (Fig. 2F-G).

Finally, long-term examination starting from very early developmental stages shows that rupture always occurs in the same position in the amnion, but not in the serosa. During their formation, the amnion and serosa share a tissue boundary until they separate into discrete membranes at the serosal window closure stage (Fig. 5A; Handel et al., 2000; Benton et al., 2013). As the EE tissues separate, serosal cells rapidly acquire fixed positions under the vitelline membrane (Koelzer et al., 2014), while internally the embryo and amnion are not so tethered. Indeed, we find that over half of all embryos examined (59%, *n*= 69) rotate longitudinally during early germband extension relative to the serosa: up to 90° about the anterior-posterior axis, with a tendency to rest laterally (Fig. 5B, Figure 5-figure supplement 1). Rotation occurs irrespective of whether the embryo’s long axis is orthogonal (this study) or parallel (Strobl and Stelzer, 2014) to gravity(Strobl and Stelzer, 2014), suggesting an inherent anisotropy of the early egg. However, there is no counterpart in later development (*n*= 118), including for 15 embryos filmed continuously and which exhibited typical frequencies of early rotation. This is consistent with the timing of amnion-serosa adhesion to form the bilayer. Thus, early rotation without any later reversal is a feature of wild type *Tribolium* embryogenesis, and apposition of the mature amnion and serosa need not correspond to where they had initially detached from one another.

**Figure 4.**
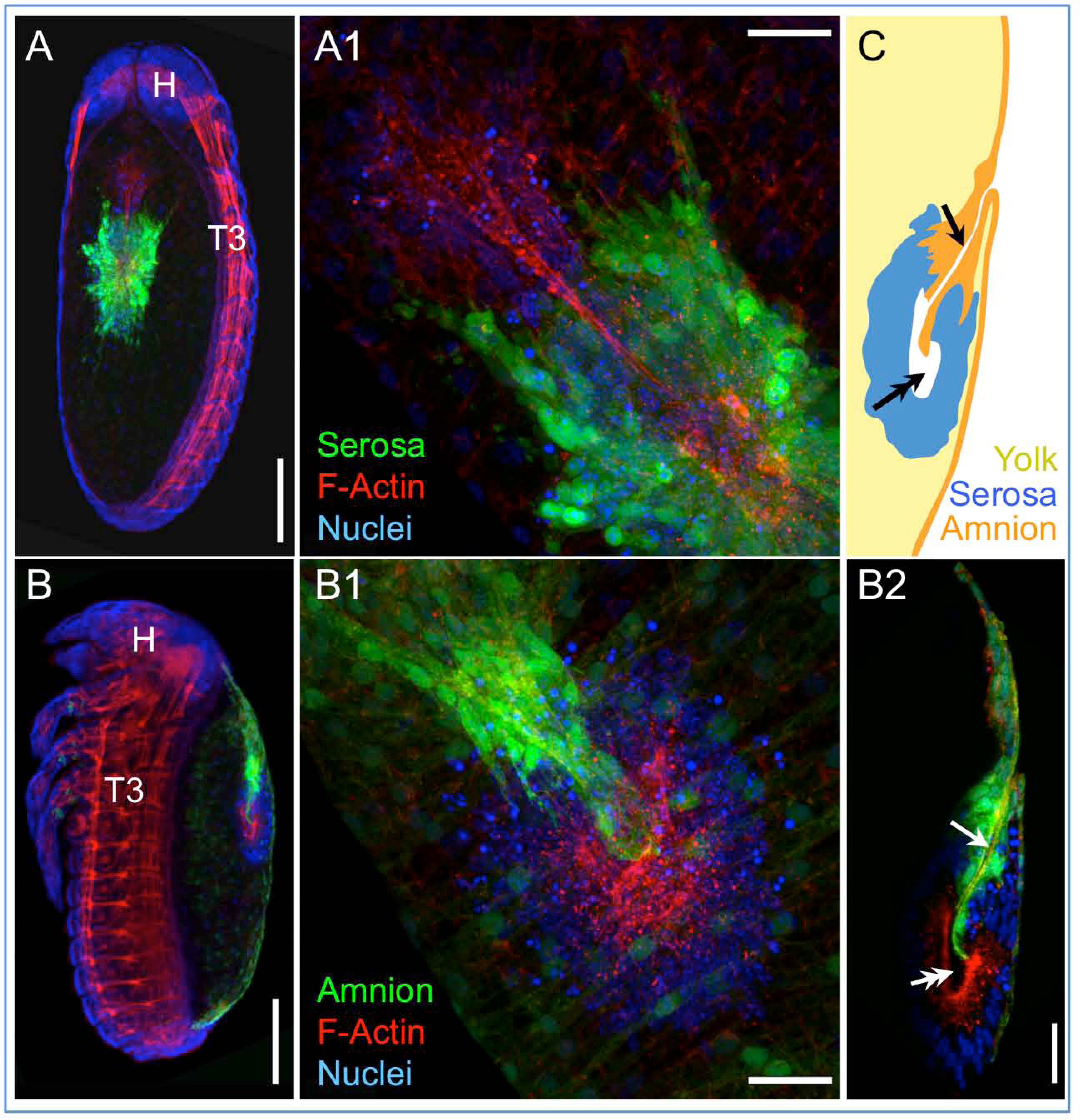
Separation during degeneration: the amnion and serosa resolve into two distinct dorsal organ structures. Images are dorsal (A,A1,B1) or lateral (B,B2,C), with anterior up (A,B,B2,C), or upper-left (A1,B1), shown as maximum intensity projections (A,A1,B,B1) or sagittal optical sections (B2,C). Tissue-specific EGFP labels two distinct dorsal organ structures, each characterized by apical F-actin enrichment around a hollow center or cleft (arrow: amniotic; double-headed arrow: serosal). Scale bars are 100 *μ*m (A,B) and 25 *μ*m (A1,B1,B2). After the stage shown here, the two dorsal organs will separate: the serosa remains medial, while the amniotic dorsal organ migrates anteriorly to a position behind the head before undergoing final degeneration (Panfilio et al., 2013).

**Figure 5.**
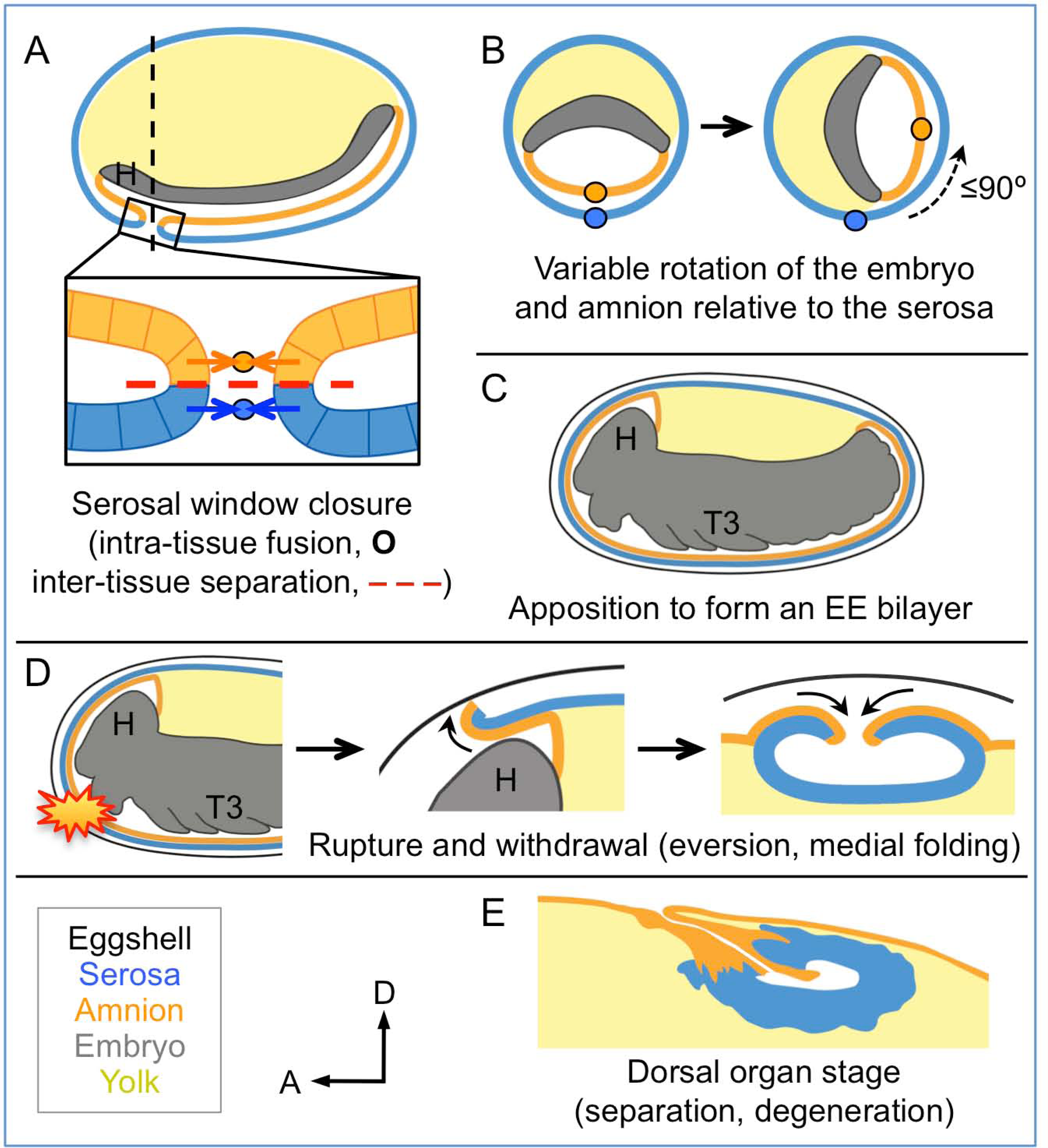
Changing amnion-serosa interactions during *Tribolium* extraembryonic morphogenesis. Schematics are sagittal, with anterior left and dorsal up (A,C-E), or transverse with dorsal up (B), illustrating: initial separation during formation of the amnion and serosa as distinct epithelial covers (A), relative rotation in many embryos (B: shown at the position of the dashed line in A, see also Figure 5-figure supplement 1), EE apposition at the retracted germband stage (C), the successive stages of EE tissue withdrawal (D), and final tissue structure during degeneration at the dorsal organ stage (E, reproduced from Fig. 4). Amnion-serosa apposition persists during withdrawal morphogenesis (D), which involves EE tissue rupture (starburst), curling over of the resulting tissue edge as it everts (shown for anterior-dorsal edge), and early serosal compaction as the EE edges fold medially. Abbreviations as in previous figures.

Given that rupture invariantly occurs in the same region in the amnion, variable longitudinal rotation of the embryo and amnion away from where they had detached from the early serosa therefore precludes an impetus for EE rupture from any serosa-or eggshell-specific landmark associated with initial EE tissue separation (*cf*., Strobl and Stelzer, 2014). Rather, later serosal regionalization (Koelzer et al., 2014) may reflect the post-rotation position of the embryo and amnion. The region of amniotic specialization for rupture may be autonomous (*e.g.*, corresponding to where it ultimately closed at the serosal window stage) or induced by signals from the embryo (the underlying head structures), but is unlikely to be determined by the serosa. In future work, identifying the molecular cues for amniotic regionalization and the proximal triggers for rupture itself will further clarify the amnion’s role.

### Serosal contractility drives withdrawal morphogenesis

Once the EE tissues have ruptured anterior-ventrally, they pull back from the embryo as an everting sack that folds up to a dorsal-medial position, similar to a pillowcase being turned inside out as it is peeled off of a pillow (Fig. 5D). The serosa appears to be the driving force for withdrawal. We had previously observed that the serosa facilitates the final stages of withdrawal during dorsal closure, making the process more robust and efficient (Panfilio et al., 2013). Here, we find that this is true throughout these morphogenetic movements.

Firstly, we again use the serosa-less situation after *Tc-zen1*^*RNAi*^ to assess the serosa’s normal contribution. Wild type withdrawal is rapid, with visible squeezing of the embryo’s abdomen as the posterior EE tissue edge pulls back and clears the posterior pole (Fig. 3E), due to serosa-specific enrichment in apical F-actin as the cells undergo a drastic shape change to become pseudocolumnar (Figs. 2F-G, 2I-J, 3H). Indeed, wild type specimens occasionally exhibited a transient extra fold in the posterior EE tissue (Fig. 2G,G′), likely reflecting a degree of stochasticity in tissue relaxation after snapping over the abdomen, similar to kinetic descriptions in other species (e.g., Patten, 1884). Both the histological and kinetic situations are altered in the absence of the serosa. In *Tc-zen1*^*RNAi*^ embryos the amnion does withdraw dorsally, but over a much longer time scale without sufficient force to squeeze the abdomen away from the eggshell, with tissue ruffling into folds rather than eversion due to apical constriction, and without significant F-actin enrichment or increase in cell height (Fig. 3F,G,I).

As a second approach to functionally test the serosa’s role, we then used RNAi against *Tc-zen2*, the paralogue of *Tc-zen1*. *Tc-zen2* is expressed extraembryonically, and strong RNAi knockdown completely blocks EE rupture (van der Zee et al., 2005). We find that the *Tc-zen2*^*RNAi*^ serosa still contracts strongly (Fig. 3J-K). Although the amniotic cavity remains unopened and there are no free EE tissue edges, the serosa contracts until it tears ectopically, withdrawing ventrally towards the amnion, to which it remains attached throughout the bilayer region. Extensive amnion-serosa attachment can also be inferred from weaker *Tc-zen2*^*RNAi*^ phenotypes, when tearing of the EE tissue produces a constrictive EE “belt” around the embryo (Fig. 3L-M). Whether visualized for the serosa or amnion, the nature of embryonic constriction is consistent with the intact portions of both tissues contracting together. Thus, we propose that correct rupture of the amnion is required for directed EE tissue withdrawal, while the serosa autonomously provides the motive force to achieve this.

### The EE bilayer resolves into two dorsal organs during dorsal closure

The dorsal organ is a transient, hollow structure formed by the serosa as it sinks into the yolk and degenerates in many insects (Panfilio, 2008). Importantly, it is represented as comprising only serosal tissue, with the amnion restricted to attachment to the serosa at its edges (Enslee and Riddiford, 1981; Tojo and Machida, 1997; Panfilio and Roth, 2010) or fully enveloping the serosa as a smooth outer layer (Rempel and Church, 1971), if a distinct amnion was recognized (see first results section). The persistent bilayered structure of the *Tribolium* EE tissues, however, results in both a serosal dorsal organ and a nested, amniotic dorsal organ (Fig. 4). In light of these observations, re-analysis of late serosa-less *Tc-zen1*^*RNAi*^ embryos indicates that the previously observed region of anterior-medial amniotic F-actin enrichment (Panfilio et al., 2013) corresponds to the amniotic dorsal organ, demonstrating that this structure also forms in the absence of a serosa. Moreover, in wild type at the end of the EE tissues’ lifetimes, they once again function as independent structures, involving separation of the bilayer as the tissues degenerate separately (Fig. 4).

## CONCLUSIONS

Altogether, the tissue reorganizations for insect EE withdrawal have complex implications for inter-epithelial attachment, balancing a requirement for tissue continuity over the yolk and coordinated withdrawal with enabling the amnion and the serosa to follow their own morphogenetic programs. It was previously known that the *Tribolium* EE tissues arise from the same blastodermal cell sheet before enclosing the embryo and detaching from one another (Handel et al., 2000; Benton et al., 2013; Koelzer et al., 2014) (Fig. 5A). After those early stages, the structure and arrangement of the amnion had been obscure. Here, we have characterized a new genetic tool that literally illuminates this enigmatic tissue for the first time. Following its early detachment from the serosa, the amnion often rotates relative to the serosa (Fig. 5B) before reattaching to form the bilayer (Fig. 5C). Then, given that the EE tissues withdraw from the embryo as a single unit (Fig. 5D), the re-establishment of their independence during final degeneration at the dorsal organ stage is striking (Fig. 5E). The mechanical requirements of these morphogenetic events imply a precisely regulated and dynamic mode of epithelial attachment. The apposed surfaces (Fig. 5C-D) present a basal-basal interface where they presumably interact via a basement membrane, which is known to influence both cell adhesion and shape changes at tissue folds during animal embryogenesis (Daley and Yamada, 2013). The withdrawing EE tissues of *Tribolium* thus provide a new system for investigating rapid remodeling of basement membrane structure and cellular-extracellular matrix interactions in simple epithelia.

At the same time, the bilayered EE arrangement in *Tribolium* is thus far unique among insects. While limited data are available for holometabolous insects with complete EE tissues, hemimetabolous insects have been more extensively studied and have a different arrangement. Hemimetabolous EE withdrawal, known as katatrepsis, involves the embryo being pulled out of the yolk by the EE tissues. The requirements for katatrepsis restrict amnion-serosa connection to a distinct border region with lateral-lateral cell contact (Panfilio, 2009; Panfilio and Roth, 2010), but katatrepsis was lost at the base of the holometabolous insect radiation (Panfilio, 2008). Further taxonomic sampling will reveal whether *Tribolium* represents the norm or one of multiple possible EE configurations within the Holometabola, where non-katatreptic withdrawal may have relaxed morphogenetic constraints on extraembryonic developmental strategies.

## MATERIALS AND METHODS

### *Tribolium* stocks and RNAi

Analyses were performed in the San Bernardino wild type strain, EFA-nuclear-GFP (nGFP) line (Sarrazin et al., 2012), and selected GEKU screen enhancer trap lines (Trauner et al., 2009): G12424 (“serosa”) and G04609 (“heart”: cardioblasts and segmental domains), as described (Koelzer et al., 2014); and HC079 (“amnion”, characterized in this study, see also Figure 1-figure supplement 1, Movie S1). RNA interference (RNAi) for *Tc-zen1* and *Tc-zen2* was performed as previously described, with double-stranded RNA of ≥688 bp at 1 *μ*g/*μ*l concentration (van der Zee et al., 2005; Panfilio et al., 2013).

### Live imaging

Embryos were dechorionated and mounted on slides in halocarbon oil for conventional microscopy (Panfilio et al., 2013). Additionally, a light sheet fluorescence microscope (mDSLM model: Strobl and Stelzer, 2014) was used for high temporal resolution imaging. Here, embryos were dechorionated and embedded in low melt agarose and filmed in anterior-ventral aspect (see Supplementary file 1 for acquisition details). Subsequent hatching was confirmed in all cases. Cell tracking was performed on maximum intensity projection (MIP) time-lapse movies with MTrackJ (Meijering et al., 2012).

### Rupture initiation analysis

Embryos were recorded every 3.5 minutes at 27.5-28°C on a DeltaVision RT microscope (Applied Precision), with simultaneous recording of up to 11 embryos of both the serosa-heart and amnion-heart crosses in each of three experiments. Using the MIP movie output, rupture initiation was determined as the interval between the following events. The start of rupture was recorded as either the first frame in which two or more cells clearly diverged, followed by the appearance of a hole in the EE tissue at the same position in subsequent frames, or as the first frame after a sudden collapse of the anterior EE membrane, whichever came first. The end of rupture was recorded as the first frame in which the retracting membrane cleared the head dorsally.

### Longitudinal rotation analysis

Embryos were analyzed from time-lapse data sets of at least 15.75 hours’ duration at 30°C (see also Figure 5-figure supplement 1). Orientation and degree of rotation were determined by eye from MIP movies, and categorized into eight sectors of 45° around the egg circumference.

### Histological staining

To assess early amnion topography, fuchsin staining after standard fixation and methanol shock without devitellinization was performed based on standard protocols (Wigand et al., 1998). For fluorescent imaging of endogenous EGFP, embryos were fixed, manually dechorionated, optionally stained with fluorescently conjugated phalloidin or wheat germ agglutinin (WGA), and embedded in Vectashield mountant with DAPI as described previously (Panfilio et al., 2013).

## SUPPLEMENTAL INFORMATION

Supplemental Information includes 4 figures, 3 movies, and 1 supplementary file (table).

## AUTHOR CONTRIBUTIONS

K.A.P. conceived the experiments. S.K. and K.A.P. planned the RNAi and fuchsin experiments; T.H., M.H., and K.A.P. designed the rupture initiation experiments; M.H. designed the light microscopy experiments. All authors conducted the experiments, analyzed the data, and helped draft or revise the manuscript.

## ACKNOWLEDGMENTS

We thank Sue Brown for providing many enhancer trap lines; Gianluca Sharbaf Azari, Max Kornilov, and Max Pentzien for contributions during student research; Frederic Strobl, Sven Plath, Ernst Stelzer, and Pavel Tomancak for training and advice on light sheet microscopy; Ferdinand Grawe (Microscopical Anatomy and Molecular Cell Biology Research Group, Institute I for Anatomy, University of Cologne) for assistance with electron microscopy. This work was supported by the German Research Foundation (Deutsche Forschungsgemeinschaft) Emmy Noether Program (grant number PA 2044/1-1 to K.A.P.).

## COMPETING INTERESTS

The authors declare that they have no competing financial or non-financial interests.

## FIGURE SUPPLEMENT AND SUPPLEMENTARY MOVIE LEGENDS

**Figure 1-figure supplement 1. Developmental time course of EGFP expression in the enhancer trap line HC079.**

**A-B.** The enhancer trap line HC079 was mapped to a gene-poor, intergenic region on chromosome 3. EGFP signal is first detected in the amnion during germband retraction, becoming increasingly bright throughout the tissue up to the time of serosal rupture, which heralds the onset of withdrawal morphogenesis. Relative EGFP signal quantification is plotted as the mean ± standard deviation during a 48-hour time-lapse recording, with corresponding values for landmark developmental stages (GBE, germband extension; GBR, germband retraction; SR, serosal rupture; MT, muscle twitches). See Koelzer, *et al*., (2014) for mapping technique, original definitions, acquisition parameters, and analysis method. Note that “serosal rupture” is the staging term but in fact refers to first discernible rupture of both EE tissues at the beginning of withdrawal morphogenesis. Landmark stage values are also based on nGFP embryos that were recorded in parallel for calibration (combined n= 7-23 embryos per stage). Vertical red lines indicate the mean SR age. In A, relative EGFP signal is the mean grey value (n= 10), shown for minimum age (from a four-hour egg collection) relative to serosal rupture. In B, signal is the integrated density of the posterior and anterior egg halves (0-50% *vs*. 50-100% egg length) for embryos recorded in both lateral and ventral aspect (n= 5), with standard deviation shown on only one side of each plot for clarity. Here, the x-axis shows minimum absolute age in hours after egg lay (hours AEL). Vertical black lines and asterisks indicate the first time points at which the anterior and posterior EGFP values differ significantly (*: p<0.05, **: p<0.01 for paired, two-tailed *t*-tests). Although this difference in part reflects the relative area of visible amnion in lateral aspect (greater anteriorly), the trend still holds when only embryos in ventral aspect are considered (uniform relative amnion surface area).

**C.** In our examination of rupture initiation in the amnion (HC079) × heart (G04609) cross compared to the serosa (G12424) × heart (G04609) cross, we found that the amnion cross develops at a faster rate, shown here as the minimum age at the time of serosal rupture (x-axis; amnion cross: 3007 (median) ± 81 (MAD) minutes, n= 28; serosal cross: 3168 (median) ± 85 (MAD) minutes, n= 30; *p*= 4.942e-06, Mann-Whitney U test), shown in relation to the duration of rupture initiation (y-axis: same as main text Fig. 3C). However, even when a normalization factor was introduced to adjust for developmental rate, rupture initiation is longer in the amnion background. **D-I.** Selected still images of an example embryo in lateral aspect (deconvolved maximum intensity projections; anterior left, dorsal up), showing EGFP signal predominantly in the amnion as well as in the eye (E) and multiple ringed domains within the legs (arrows). After dorsal closure (I), weak EGFP signal is also seen segmentally and in the usual nervous system tissues in which the core 3xP3 promoter drives expression (Koelzer et al., 2014). Scale bar (shown in D) is 100 *μ*m. The double-headed arrow in D marks the posterior tip of the abdomen, indicating that the embryo is midway through germband retraction at this time. See also Movie S1.

**Figure 2-figure supplement 1. The amnion and serosa form a persistent bilayer comprised of two morphologically distinct epithelia, including in the anterior-ventral rupture competence zone.** Although EGFP intensity varies between cells, EGFP signal for a single EE tissue type labels all cells within a continuous epithelial sheet. Moreover, the amnion and serosa have strikingly different cellular morphologies, enabling cells of both tissues to be distinguished, with the serosa overlying the amnion in maximum intensity projections.

**A-C.** The two EE tissues can be readily distinguished by their cellular morphology, and both can be discerned as continuous epithelia within the rupture competence zone (shown after the completion of germband retraction, at 44-48 hours after egg lay). Shown are two examples with serosal EGFP (A-B) and one with amniotic EGFP (C; see also main text Fig. 2D). “Overview” images show lateral, ventral, and ventral-lateral views of maximum intensity projections (A1,B1,C1, respectively) with anterior left and the region of interest around the head appendages indicated by the blue boxed region (due to the curvature of the egg, only the regions actually visible in high-magnification projections of thinner z-stacks are indicated). “Merge” images (A2,B2,C2) show maximum intensity projections of the indicated region, labeled for F-Actin (red, phalloidin), nuclei (blue, DAPI), and the specified EE tissue (green, EGFP). Note that embryonic-specific signal (F-Actin and DAPI signal for small cells not in contact with the tissue at the egg surface) was manually deleted from deeper optical sections with the freehand selection tool in ImageJ before rendering the projected images shown here. “Schematic” images (A3,B3,C3) show cell and nuclear outlines. Cell outlines were determined by EGFP signal (for A3,B3,C3) and F-Actin signal (for C3, for serosal outlines only). Nuclear outlines were determined by EGFP signal or DAPI (for all EGFP-negative nuclei only). Serosal cells are shown in cyan or blue; amniotic nuclei and cells are shown in orange or white. Dashed outlines highlight individual serosal (red) and amniotic (orange) cells, which are also labeled by arrows of the same color in the single-channel micrographs for EGFP (A4,B4,C4) and F-Actin (A5,B5,C5). Strong F-Actin signal correlates with serosal cell shape (compare A4,A5 and B4,B5), such that F-Actin signal can be used to infer serosal cell shape even when that tissue is not labeled with EGFP (C3,C5). Furthermore, the F-Actin-inferred serosal cell outlines correspond to EGFP-negative nuclei in the amnion enhancer trap line (C3; see also main text Fig. 2D2). Note that in order to preserve anterior-ventral EE tissue structure and topography, the specimens presented here (A-C) are still within the vitelline membrane. After fixation, embryos were cut in half (transversely) with a razor blade to allow the phalloidin and DAPI staining reagents to penetrate into the sample. (The EGFP is the endogenous signal.)

**D-E.** This bilayer arrangement persists for over a day prior to rupture, with no cell mixing or change in cell number in either the serosa (D) or the amnion (E). Selected stills from time-lapse movies are shown at the times indicated relative to rupture (at 30°C), in anterior-ventral aspect and oriented with the ventral region uppermost, with light sheet illumination coming from above. Individual nuclei were tracked hourly, showing the static nature of serosal cells until the time of rupture, while amniotic nuclei move much more within the cells and the amniotic tissue itself shifts slightly anterior-dorsally. The initial opening at the time of rupture is indicated with an asterisk (D4,E4). Axes were determined based on weak embryonic signal (D5,E5; the withdrawing edge of the amnion is still visible in E5). Embryos were recorded with a light sheet microscope every 2 minutes. Images shown here are maximum intensity projections with gamma corrections 0.5 (serosa) and 0.7 (amnion) as well as brightness correction to accommodate for changes in signal intensity of these lines during development (see Figure 1-figure supplement 1A and Koelzer et al., 2014).

**F.** The bilayer arrangement can also be seen in transmission electron micrographs of sagittal sections (examined after the completion of germband retraction, at 44-48 hours after egg lay; see also main text Fig. 2B-C). The red box on the semithin section (1*μ*m thick, stained with toluidine blue) approximately indicates the anterior-ventral region shown in F2-F3. The arrow highlights an intercellular junction within the serosa (F2), and false coloring highlights the continuity of the two distinct epithelial layers (F3). Both EE tissues have very flat cells, with an apical-basal thickness of only ∼200 nm in the amnion and ∼450 nm in the serosa, in the aponuclear region shown here. Ultrathin sections (50-100 nm thick) were cut with an “Ultra” diamond knife on a Leica Ultracut UCT microtome, and examined with a Zeiss EM 109 electron microscope. Scale bars are 50 *μ*m (A1,B1,C1,D1-D5,E1-E5), 20 *μ*m (A2-A5,B2-B5,C2-C5), and 1 *μ*m (F2-F3). Abbreviations: An, antenna; Am, amnion; D, dorsal; H, head; L, left; Lr, labrum; M, mandible; Mx, maxilla; R, right; Ser, serosa; T2, second thoracic segment; V, ventral ; Vm, vitelline membrane.

**Figure 3-figure supplement 1. Comparison of early tissue opening in the amnion and serosa.**

**A-B.** The embryos are heterozygotes expressing EGFP in the specified extraembryonic tissue and in the embryonic landmark domains of the “heart” enhancer trap line. They are shown in anterior-ventral views as maximum intensity projections of single time points taken from z-stacks acquired every 20 seconds with a light sheet microscope. Light sheet illumination is coming from above in both recordings. Image A is reproduced from main text Figure 3 for side-by-side comparison with image B. The colored time scale shows the duration of track segments over 36.7 minutes at 19.5±1°C. The embryos are stage matched for the degree of EE tissue withdrawal by the end of the 36.7-minute interval, when the EE tissue has cleared the proximal portion of the first leg pair (T1). Along the time scale and superimposed on the embryo are the site of rupture (white asterisk) and the withdrawing EE tissue edge (white lines, line with arrowhead marks time point shown). In the main text, statistical analysis of the duration of “rupture initiation” was based on embryos viewed in lateral aspect and was defined as the time from first discernible opening to when the EE tissues cleared the embryo’s head dorsally (see Methods). Since dorsal head structures are not visible in the anterior-ventral movies shown here, the approximate equivalent duration is through the time when the maxillae are cleared (Mx): the interval from 9:00 to 20:20 in the amnion (11:20 duration) and from 20:00 to 27:20 in the serosa (7:20 duration). Tracks are shown for selected nuclei. Scale bars are 50 *μ*m. See also Movie S3.

**Figure 5-figure supplement 1. Frequency of embryonic longitudinal rotation during development.**

**A.** Sample sizes and film durations (blue bars) relative to landmark developmental stages (yellow bars: SW, serosal window closure; GBE, germband extension complete; SR, serosal rupture; staging as in Koelzer, *et al*. (2014)). Embryos were analyzed from multiple time-lapse data sets of at least 15.75 hours’ duration at 30°C, recorded with an inverted DeltaVision RT (Applied Precision) microscope. Pooled data included embryos in the nGFP and various enhancer trap backgrounds (G04609-heart, G12424-serosa, KT650-serosa, HC079-amnion), including heterozygote crosses of HC079 with each of G04609, G12424, and nGFP. (Background autofluorescence was sufficient for scoring early development in the absence of specific GFP signal.) “Frequency by stage” refers to the beginning of rotation; all embryos beginning during (16%) or after (41%) SW completed rotation before GBE. Rarely, a moderate degree of rotation occurred during dorsal closure (4%).

**B.** Chi-square statistics for specified comparisons. For significant results, text color-coding indicates the nature of the interaction. Longitudinal rotation ranged from approximately 20 to 100 degrees, with two-thirds of embryos rotating approximately 90°. Embryos laying on their dorsal or ventral surface (“poles”) were more likely (69%) to rotate 90°, while embryos in lateral aspect (85%) tended to only rotate 45°. While overall there was no bias in direction of rotation, embryos initially laying on their right sides tended to rotate anticlockwise (70%) while embryos on their left sides rotated clockwise (78%). Direction of rotation was from the perspective of the embryo’s posterior (*i.e.*, clockwise is right, anticlockwise is left). These directional biases are applicable across the full range from ventral-lateral through dorsal-lateral, such that rotation tended to result in more strictly lateral final orientations (64%). Meanwhile, most non-rotating embryos began in a lateral orientation (89%).

**C.** Pie charts representing the eight 45°-sectors of the egg circumference (D, dorsal; DL-L and -R, dorsal-lateral-left and -right; L-L and -R, lateral-left and -right; VL-L and -R, ventral-lateral-left and -right; V, ventral), color-coded for frequency of occurrence according to the heat map.

**Movie S1. Time course of EGFP expression in the *Tribolium* enhancer trap line HC079.** The embryo is shown in lateral aspect with anterior left and dorsal up. Strengthening EGFP signal is specific to the amnion during germband retraction and up to the point of EE tissue rupture and withdrawal morphogenesis. In the later stages, additional embryonic expression domains occur in the eye and legs. Throughout the movie, wandering yolk globules also exhibit low levels of EGFP, a feature observed for other enhancer trap lines from this screen (Koelzer et al., 2014). The time stamp specifies minimum age from a four-hour egg collection, in hours after egg lay (AEL). Deconvolved maximum intensity projections from a z-stack (5 *μ*m step size for a 60 *μ*m stack) recorded every ten minutes over a 48-hour time-lapse at 30°C, acquired with an inverted DeltaVision RT microscope (Applied Precision). Scale bar is 100 *μ*m. Selected still images are shown and further described in Figure 1-figure supplement 1.

**Movie S2. *Tribolium* extraembryonic tissue rupture and withdrawal shown with amnion-specific EGFP (enhancer trap line HC079).**

The embryo is shown in lateral aspect with anterior left and dorsal up, expressing EGFP from an enhancer trap with amnion-specific expression, as well as restricted late embryonic expression domains in the legs and body segments (see also Figure 1-figure supplement 1). The movie spans late amnion morphogenesis, from 3.3 hours before rupture through late dorsal closure. Of particular note are the brighter, rounder amniotic cells in an anterior-ventral cap, in which rupture occurs. During withdrawal, the amnion everts (turns inside out), such that the surface that had faced inward toward the embryo is flipped outward to face the vitelline membrane: this is particularly apparent from 35 to 49 minutes after rupture as the ruptured tissue edge folds over. Time is shown relative to tissue rupture at 0 minutes. Images are maximum intensity projections with a gamma correction of 0.7 from a z-stack (7 *μ*m step size for a 77 *μ*m stack) recorded every seven minutes over a 9.9-hour time-lapse at 24°C, acquired with an Axio Imager.Z2 with ApoTome.2 structured illumination (Zeiss). Scale bar is 100 *μ*m. Main text Fig. 3B shows a still at 21 minutes after rupture.

**Movie S3. Extraembryonic rupture filmed at high temporal resolution.** Embryos are shown in anterior-ventral aspect with anterior left, and labeled with EGFP from a heterozygote cross of enhancer trap lines labeling the amnion (line HC079) or serosa (line G12424) with selected embryonic domains in the head, segments, and legs (line G04609, “heart”). During the movie, the EE tissues rupture and withdraw to the extent that the head and first leg pair are exposed. Selected EE nuclei were tracked, where track color segments correspond to fixed units of time (red, orange, yellow, and green: 8.3 minutes; blue: final 3.3 minutes). Elapsed time is shown. Images are maximum intensity projections from a z-stack (2.58 *μ*m step size for a 175 or 200 *μ*m stack) recorded every 20 seconds at 19.5±1°C and shown for a 36.7-minute period (see Supplementary file 1 for further acquisition details). Single-sided light sheet illumination is from the top and recorded in one angle of view to maximize acquisition speed, resulting in deterioration of the signal in the lower part of the image due to scattering. The final frame shows time point 36:40 with the approximate position of mouthparts labeled for orientation: An, antennae; Lr, labrum; Mx, maxillae; Lb, labium; T1, first thoracic segment. Scale bar is 50 *μ*m. See also see also Figure 3-figure supplement 1 for a visual summary.

## SUPPLEMENTARY FILE: TITLE

**Supplementary file 1. Acquisition parameters for mDSLM light sheet experiments.**

## REFERENCES

Aldaz, S., Escudero, L. M. and Freeman, M. (2010) ‘Live imaging of *Drosophila* imaginal disc development’, Proc. Natl Acad. Sci. USA 107(32): 14217–14222.

Benton, M. A., Akam, M. and Pavlopoulos, A. (2013) ‘Cell and tissue dynamics during *Tribolium castaneum* embryogenesis revealed by versatile fluorescence labeling approaches’, Development 140: 3210–3220.

Calvin, S. E. and Oyen, M. L. (2007) ‘Microstructure and mechanics of the chorioamnion membrane with an emphasis on fracture properties’, Ann. N.Y. Acad. Sci. 1101: 166–185.

Daley, W. P. and Yamada, K. M. (2013) ‘ECM-modulated cellular dynamics as a driving force for tissue morphogenesis’, Curr. Op. Genet. Devel. 23(4): 408– 414.

Enslee, E. C. and Riddiford, L. M. (1981) ‘Blastokinesis in embryos of the bug, *Pyrrhocoris apterus*. A light and electron microscopic study 1. Normal blastokinesis’, J. Embryol. Exp. Morph. 61: 35–49.

Handel, K., Basal, A., Fan, X. and Roth, S. (2005) ‘*Tribolium castaneum twist*: gastrulation and mesoderm formation in a short-germ beetle’, Dev. Genes Evol. 215(1): 13–31.

Handel, K., Grünfelder, C. G., Roth, S. and Sander, K. (2000) ‘*Tribolium* embryogenesis: a SEM study of cell shapes and movements from blastoderm to serosal closure’, Dev. Genes Evol. 210(4): 167–179.

Jacobs, C. G. C., Rezende, G. L., Lamers, G. E. M. and van der Zee, M. (2013) ‘The extraembryonic serosa protects the insect egg against desiccation’, Proc. R. Soc. B 280(1764): 20131082.

Jacobs, C. G. C., Spaink, H. P. and Zee, M. v. d. (2014) ‘The extraembryonic serosa is a frontier epithelium providing the insect egg with a full-range innate immune response’, Elife 3: e04111.

Kobayashi, Y. and Ando, H. (1990) ‘Early embryonic development and external features of developing embryos of the caddisfly, *Nemotaulius admorsus* (Trichoptera: Limnephilidae)’, J. Morph. 203(1): 69–85.

Koelzer, S., Kölsch, Y. and Panfilio, K. A. (2014) ‘Visualizing late insect embryogenesis: Extraembryonic and mesodermal enhancer trap expression in the beetle *Tribolium castaneum*’, PloS ONE 9(7): e103967.

Meijering, E., Dzyubachyk, O. and Smal, I. (2012) Chapter 9: Methods for cell and particle tracking. in P. M. Conn (ed.) Imaging and Spectroscopic Analysis of Living Cells — Optical and Spectroscopic Techniques vol. 504: Elsevier.

Panfilio, K. A. (2008) ‘Extraembryonic development in insects and the acrobatics of blastokinesis’, Dev. Biol. 313(2): 471–491.

Panfilio, K. A. (2009) ‘Late extraembryonic development and its *zen-RNAi*-induced failure in the milkweed bug *Oncopeltus fasciatus*’, Dev. Biol. 333(2): 297–311.

Panfilio, K. A., Oberhofer, G. and Roth, S. (2013) ‘High plasticity in epithelial morphogenesis during insect dorsal closure.’, Biol. Open 2(11): 1108–1118.

Panfilio, K. A. and Roth, S. (2010) ‘Epithelial reorganization events during late extraembryonic development in a hemimetabolous insect’, Dev. Biol. 340(1): 100–115.

Patten, W. (1884) ‘The development of Phryganids, with a preliminary note on the development of *Blatta germanica*’, Q. J. Microsc. Sci. 24: 549–602.

Rafiqi, A. M., Park, C.-H., Kwan, C. W., Lemke, S. and Schmidt-Ott, U. (2012) ‘BMP-dependent serosa and amnion specification in the scuttle fly *Megaselia abdita*’, Development 139: 3373–3382.

Rempel, J. G. and Church, N. S. (1971) ‘The embryology of *Lytta viridana* Le Conte (Coleoptera: Meloidae). VII. Eighty-eight to 132 h: the appendages, the cephalic apodemes, and head segmentation’, Can. J. Zool. 49: 1571–1581.

Rezende, G. L., Martins, A. J., Gentile, C., Farnesi, L. C., Pelajo-Machado, M., Peixoto, A. A. and Valle, D. (2008) ‘Embryonic desiccation resistance in *Aedes aegypti*: presumptive role of the chitinized serosal cuticle’, BMC Dev. Biol. 8: 82.

Sarrazin, A. F., Peel, A. D. and Averof, M. (2012) ‘A segmentation clock with two-segment periodicity in insects’, Science 336(6079): 338–341.

Schmidt-Ott, U. (2000) ‘The amnioserosa is an apomorphic character of cyclorrhaphan flies’, Dev. Genes Evol. 210: 373–376.

Strobl, F. and Stelzer, E. H. K. (2014) ‘Non-invasive long-term fluorescence live imaging of *Tribolium castaneum* embryos’, Development 141(11): 2331–2338.

Tojo, K. and Machida, R. (1997) ‘Embryogenesis of the mayfly *Ephemera japonica* McLachlan (Insecta: Ephemeroptera, Ephemeridae), with special reference to abdominal formation’, J. Morph. 234(1): 97–107.

Trauner, J., Schinko, J., Lorenzen, M. D., Shippy, T. D., Wimmer, E. A., Beeman, R. W., Klingler, M., Bucher, G. and Brown, S. J. (2009) ‘Large-scale insertional mutagenesis of a coleopteran stored grain pest, the red flour beetle *Tribolium castaneum*, identifies embryonic lethal mutations and enhancer traps’, BMC Biol. 7: 73.

van der Zee, M., Berns, N. and Roth, S. (2005) ‘Distinct functions of the *Tribolium zerknüllt* genes in serosa specification and dorsal closure’, Curr. Biol. 15: 624– 636.

Wigand, B., Bucher, G. and Klingler, M. (1998) ‘A simple whole mount technique for looking at *Tribolium* embryos’, Tribolium Information Bulletin 38: 281–283.

